# Lipid interactions are important for the Tol-Pal complex in maintaining outer membrane lipid homeostasis

**DOI:** 10.1101/2025.06.10.658765

**Authors:** Nadege Zi-Lin Lim, Robin A Corey, Wei-Chuen Poh, Phillip J Stansfeld, Shu-Sin Chng

**Affiliations:** Department of Chemistry, National University of Singapore, Singapore 117543; National University of Singapore Graduate School for Integrative Sciences and Engineering, Singapore 117456; Department of Biochemistry, University of Oxford, Oxford OX1 3QU, United Kingdom; School of Physiology, Pharmacology and Neuroscience, University of Bristol, Bristol BS8 1TD, United Kingdom; School of Life Sciences and Department of Chemistry, University of Warwick, Coventry CV4 7AL, United Kingdom; Singapore Center for Environmental Life Sciences Engineering, National University of Singapore (SCELSE-NUS), Singapore 117456

## Abstract

Gram-negative bacteria are intrinsically resistant to many antibiotics in part due to the asymmetric architecture and barrier function of their outer membrane (OM). To establish proper lipid asymmetry, cells need to ensure an intricate balance of constituent OM components, especially lipids. In this regard, the conserved, trans-envelope Tol-Pal complex plays a primary role in maintaining OM lipid homeostasis, thus OM integrity, possibly via retrograde phospholipid transport. However, mechanistic details for this process are unknown, owing to the lack of evidence for direct lipid binding. In this study, we discover that the periplasmic protein TolB, a key component of the Tol-Pal system, associates directly with membranes in vitro, via specific interactions with cardiolipin (CL). Using coarse-grained molecular dynamics simulations, we identify a CL-binding site on TolB; a single amino acid mutation at this site abolishes in vitro membrane interaction, consequently impairing cellular Tol-Pal function in maintaining OM homeostasis in *Escherichia coli*. Curiously, we find that the functional requirement for TolB-CL interactions can be partially bypassed in cells lacking CL, suggesting compensatory effects through other lipids only when CL is absent. Our findings reveal a previously unappreciated lipid-binding role for TolB, and provide novel insights into how the Tol-Pal complex may facilitate phospholipid transport across the cell envelope. Our work will inform future strategies towards developing new antibiotics against Gram-negative bacteria.

## Introduction

Gram-negative infections pose a significant threat to global health due to existing/intrinsic and emerging/acquired resistance to antibiotics.^1^ A large part of such antibiotic resistance stems from the formidable barrier conferred by the complex cell envelope, particularly the outer membrane (OM), which encapsulates the peptidoglycan layer and the inner membrane (IM).^2^ Consequently, a great many efforts have been focused on understanding Gram-negative OM biogenesis and homeostasis, culminating in well-established mechanisms for the assembly of major OM components, including β-barrel proteins,^3^ lipoproteins,^4^ and lipopolysaccharides (LPS).^5^ Even so, little is known about the bidirectional transport of bulk phospholipids (PLs), the most basic building block of any bilayer, across the cell envelope.^6–8^ In this regard, AsmA superfamily proteins and the Tol-Pal complex have only recently been implicated in anterograde (IM-to-OM) and retrograde (OM-to-IM) PL transport, respectively.^9–13^

The Tol-Pal complex is critically important for OM stability and lipid homeostasis.^10, 11^ In the absence of the Tol-Pal complex, cells accumulate excess PLs in the OM; such lipid imbalance gives rise to OM instability and permeability defects, characterized by hypervesiculation, and severe sensitivity to antibiotics and detergents.^14–17^ In fact, *tol-pal* deletion cells exhibit slower retrograde PL transport, suggesting critical involvement in the process.^11^ The Tol-Pal complex is also involved in cell division; constituent components are enriched at mid-cell during division, and mutants display defects in OM constriction, and/or septal cell wall separation (under hypoosmotic conditions).^18, 19^ However, it has now been established that septal localization of the Tol-Pal complex is not required for OM integrity, underscoring its primary role in OM lipid homeostasis and transport.^10^

The Tol-Pal complex comprises the TolQRA and TolB-Pal sub-complexes at the IM and the OM, respectively.^20^ Pal is an OM lipoprotein that either binds the cell wall,^21–23^ thus tethering the OM, or interacts with the periplasmic protein TolB.^21, 24^ The TolQRA sub-complex harnesses the proton motive force (pmf) to transduce a force on the TolB-Pal sub-complex, through TolA-TolB interaction across the cell envelope.^20^ While such force transduction is believed to be important for localization of Pal to the division site, and is hijacked by colicins for cellular entry,^20^ how the same mechanism contributes to the main function of the Tol-Pal complex in OM lipid homeostasis is unclear.^10, 11, 18, 25^ It has been proposed that the Tol-Pal complex directly mediates retrograde transport of bulk PLs.^11^ Yet, there is no evidence of lipid binding by any of the Tol-Pal components.

In this study, we establish that the function of TolB requires direct interaction with membranes, with specificity for a particular PL species, cardiolipin (CL). While trying to characterize TolB-Pal interaction at the membrane in vitro, we serendipitously discover that TolB alone binds directly to liposomes. In addition, we find that this membrane association depends on the presence of CL, both in vitro and in silico. Furthermore, molecular dynamics (MD) simulations revealed a putative CL-binding site on TolB; we demonstrate that a specific mutation at this site disrupts TolB interactions with CL-containing membranes, as well as its cellular function. Our work underscores the functional relevance of TolB-membrane association, and provides novel insights into how the Tol-Pal complex may mediate PL transport.

## Results

### TolB interacts with membranes

We initially set out to study how TolB interacts with Pal in a native environment. The strong interaction between TolB and Pal was previously observed using purified full-length TolB and truncated Pal proteins that are entirely soluble in aqueous solution.^24^ However, full-length Pal is a lipoprotein anchored to the inner leaflet of the OM via its N-terminal triacyl modification, and it is not known how the TolB-Pal interaction may be affected at/near a lipid bilayer. Therefore, we reconstituted Pal into liposomes, and examine its interaction with TolB using isothermal calorimetry (ITC). In our proteoliposomes derived from *E. coli* polar lipids, ~80% of Pal is anchored on the surface, accessible to TolB added in the external solution (Fig S1). We first showed that control titrations of TolB into buffer or buffer into Pal proteoliposomes gave negligible background heat changes. Interestingly, we also did not observe significant signals between TolB and Pal proteoliposomes, in contrast to strong heat changes detected between TolB and soluble Pal under identical conditions (Fig 1A).^24^ This unexpected result suggests either that TolB cannot bind Pal at the membrane, or that other interactions, possibly with the bilayer, mask TolB-Pal binding.

**Fig 1:**
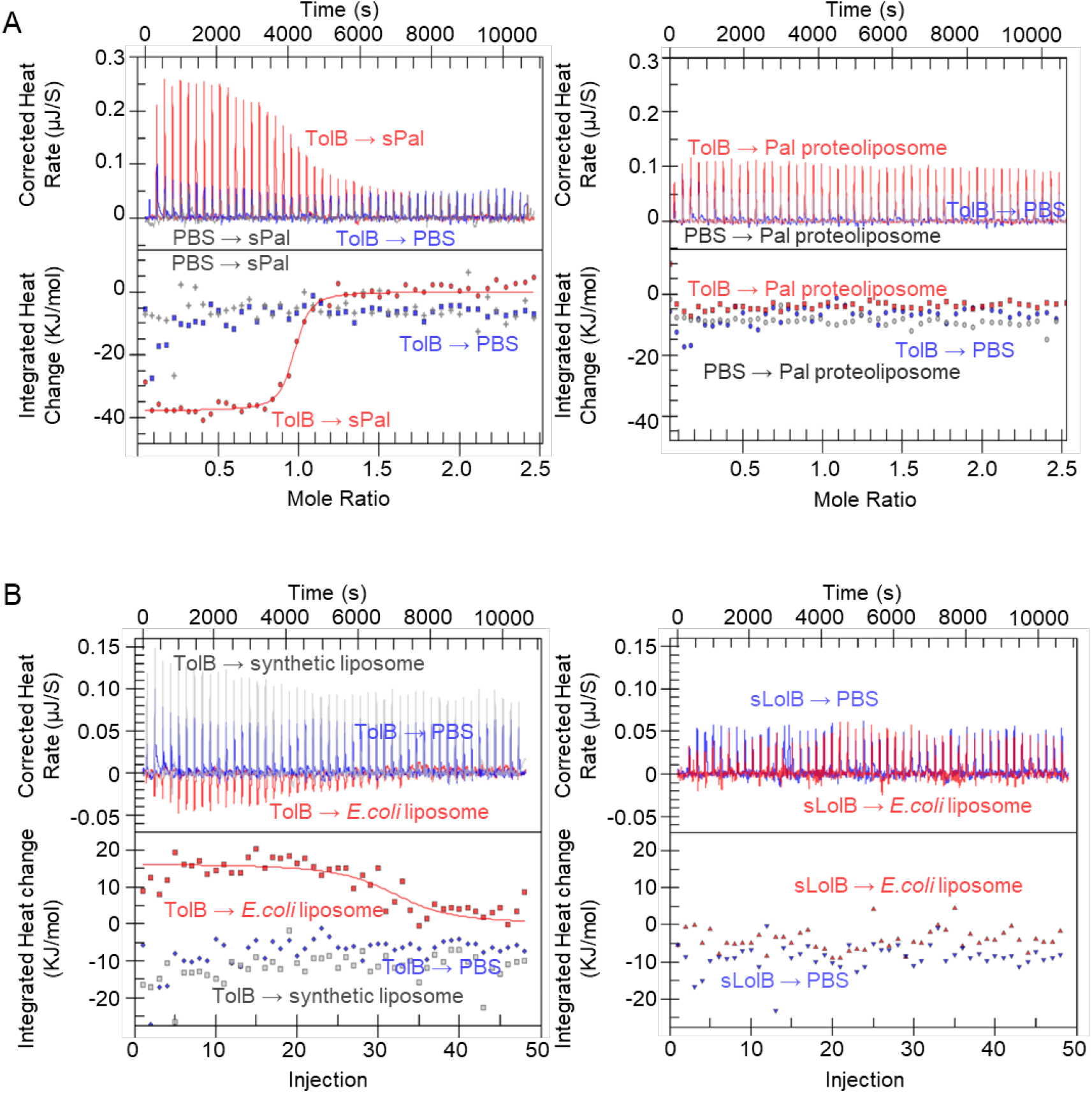
TolB interacts with lipid bilayers in vitro. (A) Representative ITC data for TolB (140 µM, 7 nmol) titrated into (*left*) soluble Pal (sPal, 20 µM, 3.32 nmol) or (*right*) Pal proteoliposomes (25 µM, 3.32 nmol surface-exposed) in phosphate-buffered saline (PBS), with indicated control titrations. (B) Representative ITC data for (*left*) TolB or (*right*) LolB (140 µM, 7 nmol) titrated into empty liposomes (5 mg lipids) composed of *E. coli* polar lipid extract (PE:PG:CL ~67:23:10) or synthetic lipids (PE:PG 3:1), with indicated control titrations. Each titration was performed at 25 oC, and repeated more than 3 times.

We hypothesized that TolB may interact with membranes, therefore titrated TolB into empty liposomes made from *E. coli* polar lipids, comprising phosphatidylethanolamine (PE): phosphatidylglycerol (PG):CL in a ratio of ~67:23:10. Remarkably, we detected endothermic heat changes between TolB and liposomes. As a control, the soluble domain of the OM lipoprotein LolB (sLolB) exhibited no interaction with empty liposomes (Fig 1B).^4^ Furthermore, we found that TolB, but not sLolB, affected the sedimentation of liposomes in an in vitro assay (Fig S2). Taken together, our results demonstrate that TolB interacts directly with the lipid bilayer.

### TolB exhibits specificity for CL binding

To understand the nature of interaction between TolB and the lipid bilayer, we also examined TolB binding to empty liposomes derived from synthetic lipids representing major species in *E. coli* (PE:PG 3:1). Interestingly, we did not observe any heat changes when TolB was titrated into these synthetic lipid liposomes (Fig 1B). TolB also did not impact these liposomes during sedimentation in vitro (Fig S2), consistent with the lack of interaction. A notable distinction between liposomes derived from synthetic lipids or *E. coli* polar lipids is the presence of CL in the latter, suggesting that TolB interacts with membranes in a CL-dependent manner.

To explore the possibility of TolB-CL interaction, we conducted interaction studies in silico using coarse-grained (CG) molecular dynamics (MD) simulations. We positioned TolB ~8 nm from membranes approximating our in vitro setup, namely PE:PG:CL at 67:23:10 (+CL), or PE:PG at 75:25 (-CL). By running 5 × 10 µs simulations in each condition, we were able to follow prospective interactions of TolB with either membrane. In the absence of CL, the distance between TolB and the bilayer fluctuated constantly, indicating brief and unstable interactions (Fig 2A). In stark contrast, when CL is present, TolB rapidly binds to the bilayer and stays bound throughout the simulations (Fig 2A). Averaging the overall time that TolB remains bound to the membrane (in all binding poses) over the 5 runs revealed an extremely high membrane binding likelihood in the presence of CL, ca. 91% vs 39% when CL was absent. These data corroborate our experimental observation that TolB interacts with membranes in a CL-dependent manner.

**Fig 2:**
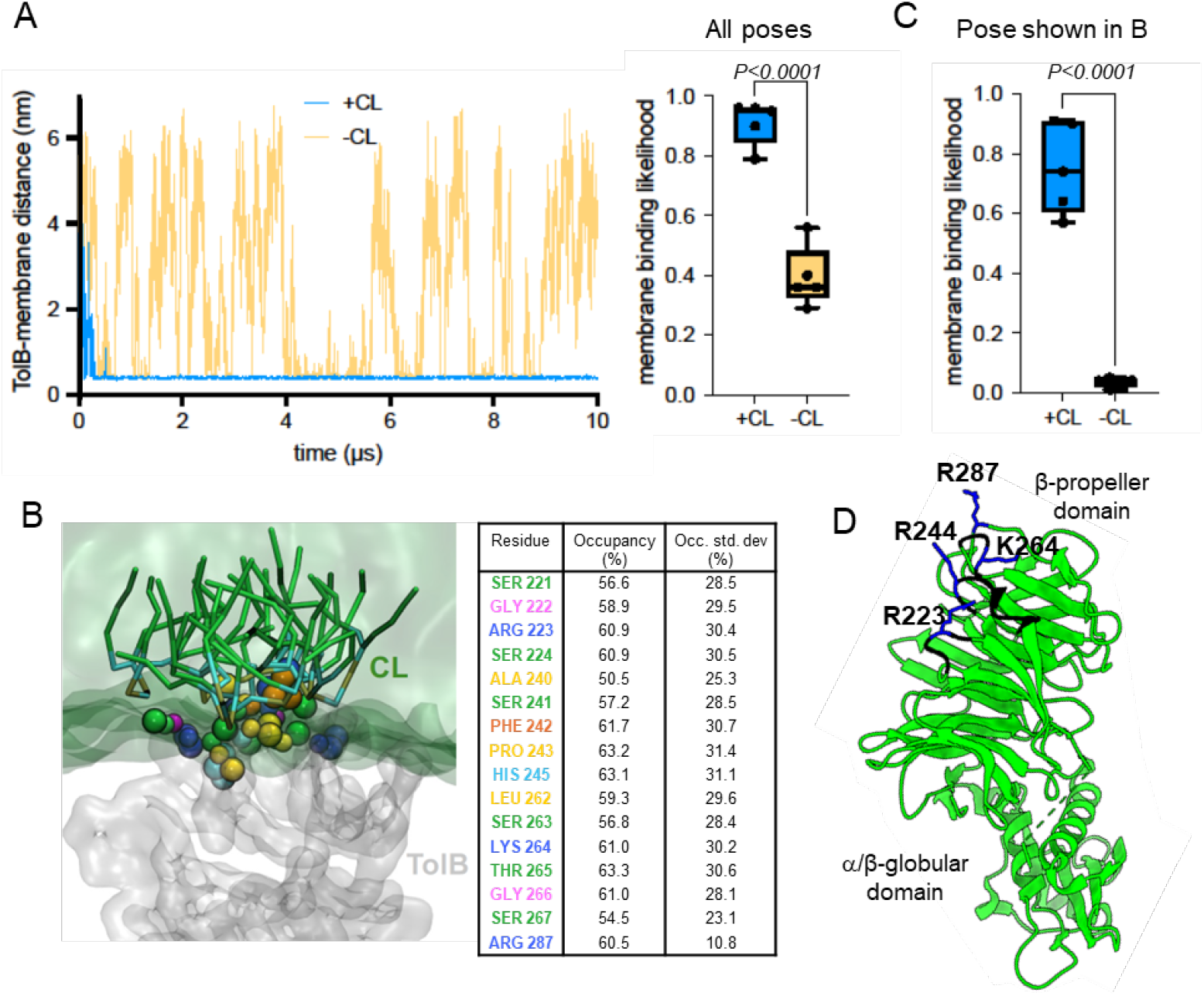
TolB interacts with CL in silico via a putative binding site. (A) Analysis of TolB-membrane distance over time in a representative 10-μs coarse-grain MD simulation, in the presence or absence of CL. Membrane binding likelihood, averaged over 5 repeat runs, considering all binding poses, is shown on the right. Two-tailed Student’s t-test was used to test for statistical significance. (B) The most representative pose of how TolB (*grey*) interacts with CL in coarse-grain simulations, showing residues (colored spheres) in contact with CL molecules (*green/cyan/tan* sticks), and their percentage CL occupancies (in table). (C) Membrane binding likelihood, just considering the binding pose in (B). (D) Contact residues are mapped onto a cartoon representation of TolB structure. Basic or other residues are colored *blue* (with side chain) or *black* (backbone only), respectively.

### Disrupting the CL binding ability of TolB abolishes its function in OM lipid homeostasis

We employed PyLipID, a software tool designed for predicting lipid binding sites within proteins, to identify possible CL binding sites on TolB based on our CG-MD data.^26, 27^ PyLipID enables the calculation of residence times and binding probabilities between proteins and lipid molecules from MD simulations. Our analysis revealed that TolB indeed harbors a CL-binding site, which remained occupied with CL for ~81% of the total simulation time, clustering on average ~4.4 CL molecules via their head groups (Fig 2B). Considering binding through this site alone, membrane binding likelihood of TolB effectively reduced to zero in the absence of CL (Fig 2C). TolB contains a C-terminal six-bladed β-propeller domain linked to a short peptide, termed the ‘TolA box’, at the N terminus, via a α/β globular domain.^28^ The TolA box and the β-propeller domain interact with TolA and Pal, respectively.^24, 28, 29^ The putative CL-binding site comprises residues from 4 adjacent loop regions in the β-propeller domain of TolB, collectively presenting 4 positively charged side chains (Arg^223^, Arg^244^, Lys^264^, Arg^287^) for interaction with the phosphate moieties in the CL head groups (Fig 2D).^27^ These Arg/Lys residues, along with neighbouring side chains, all exhibit ~60% median CL occupancy (Fig 2B).

To investigate the importance of the putative CL-binding site, we substituted each Arg/Lys position with either alanine or glutamate, and assessed whether these mutations perturbed TolB function. The alanine variants behaved similarly to wild-type TolB (Fig 3A). Remarkably, we found that R244E rendered this TolB charge-swapped variant incapable of rescuing sensitivity to SDS-EDTA and vancomycin in the Δ*tolB* strain (Fig 3A, 4A), as well as the increased cell width phenotype associated with OM lipid dyshomeostasis (Fig 4B).^10^ Importantly, this mutation did not impact TolB stability (Fig S3A), its ability to facilitate colicin A entry (Fig S3B), nor its cellular interaction with Pal (Fig S4), indicating a true and specific loss of function in maintaining OM integrity. In fact, we demonstrated that TolB^R244E^ failed to restore OM defects in a strain expressing a chimeric version of the Tol-Pal complex, whose only function is in OM lipid homeostasis along the cell periphery (Fig 3B).^10^ These findings indicate that the functional impairment caused by the R244E mutation is most likely directly linked to the disruption of TolB-CL binding.

**Fig 3:**
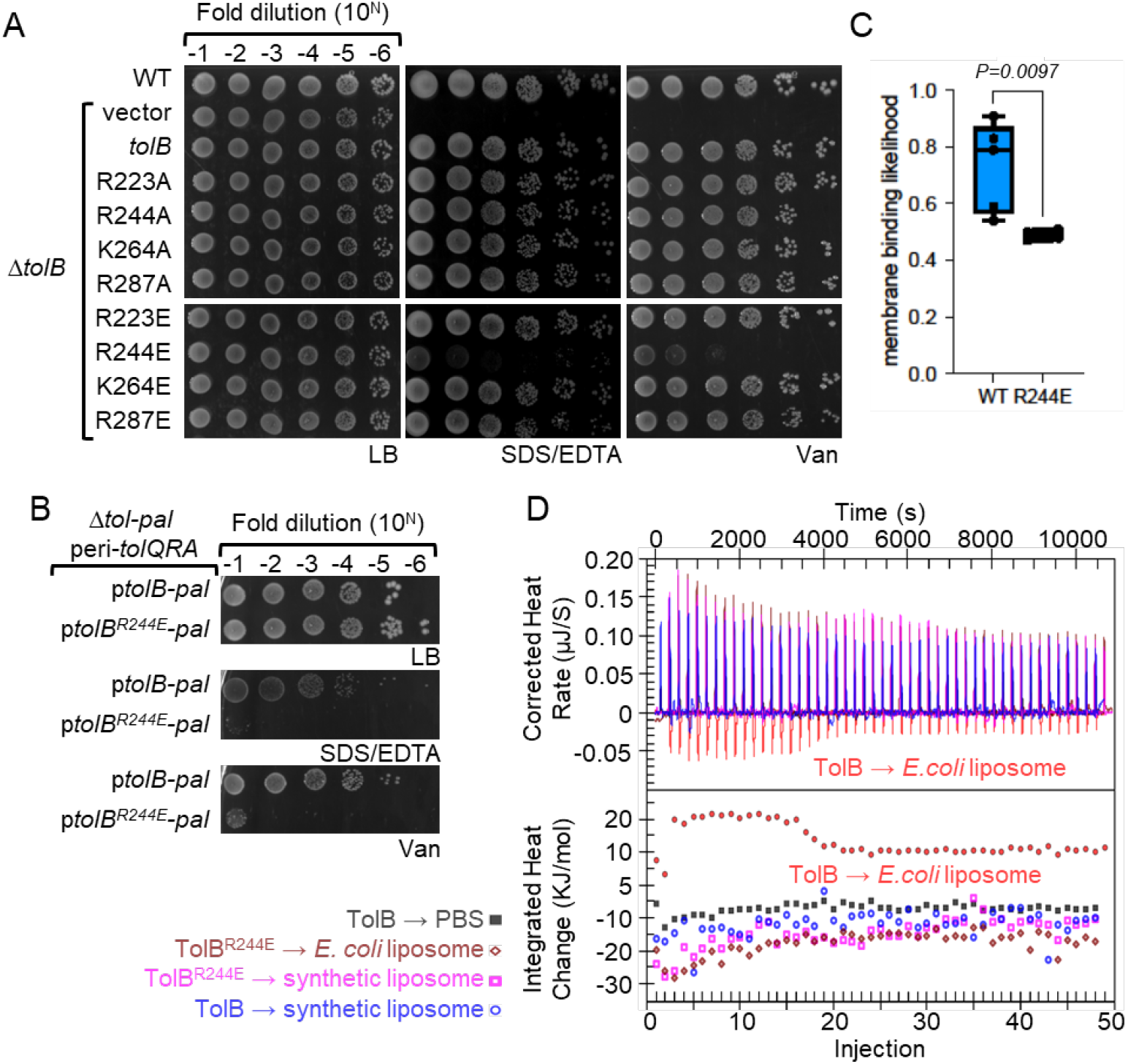
Mutation within putative CL-binding site resulted in loss of TolB function. (A) Efficiency of plating (EOP) of MG1655 wild type (WT), and Δ*tolB* strains, containing either pET22/42 empty vector or the plasmid expressing TolB and its mutants, on LB agar plates supplemented with SDS/EDTA (0.5% SDS/0.5 mM EDTA) or vancomycin (60 µg/mL) at 37oC. (B) EOP of MG1655 Δ*tol-pal* strain expressing a peripheral version of the TolQRA complex that remains randomly distributed along the cell periphery, and is known to restore OM defects,^10^ containing either pBAD43-*tolB-pal* or pBAD43-*tolB*^*R244E*^*-pal*, on LB agar plates supplemented with SDS/EDTA (0.3% SDS/0.3 mM EDTA) or vancomycin (30 µg/mL) at 37 °C. Peripheral TolQRA is engineered using a chimeric TonB-TolA that complexes with ExbBD (homolog of TolQR). (C) Membrane binding likelihood, averaged over 5 × 10 µs MD runs, of wild-type TolB and TolB^R244E^ binding to CL-containing membrane. (D) Representative ITC data of TolB and TolB^R244E^ (140 µM, 7 nmol) titrated into *E. coli* or synthetic liposomes (5 mg lipids) at 25 °C, with indicated control titrations.

**Fig 4:**
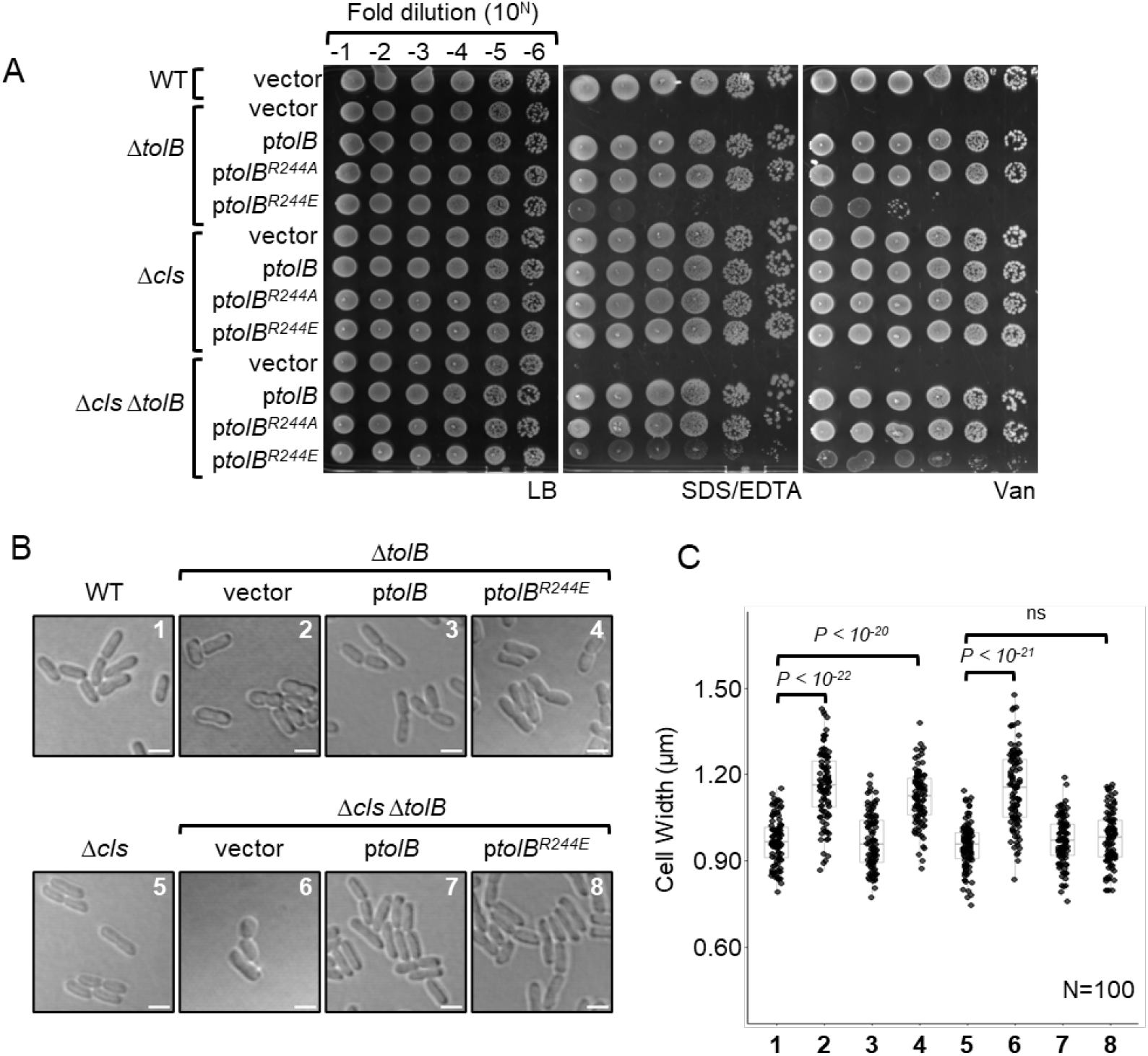
TolB^R244E^ regains partial function in strains without CL. (A) EOP of MG1655 wild type (WT), Δ*tolB*, Δ*cls*, and Δ*cls* Δ*tolB* strains, containing either pET22/42 empty vector or the plasmid expressing TolB and its mutants, on LB agar plates supplemented with SDS/EDTA (0.5% SDS/0.5 mM EDTA) or vancomycin (60 µg/mL) at 37 °C. (B) Differential interference contrast (DIC) microscope images of indicated strains from (A) grown in LB at 37 °C. Scale bar represent 2 µm. Cropped images are representative of at least three different uncropped field-of-view. (C) Quantification of the cell width of 100 individual cells of same strains (numbered) shown in (B). Box plot defines the median (center) and the 25^th^ and 75^th^ percentile (lower and upper hinge); the whisker represents the range excluding the outliers (outlier defined by sample with value further than 1.5 interquartile-range from the corresponding hinge). A two-sided Wilcoxon ranked sum test was used to test for statistical significance.

R244 occupies a central position within the predicted CL-binding site. Therefore, we hypothesize that changing this positively charged residue to a negative one (R244E) could introduce repulsive interactions with the negatively charged head groups of CL, and disrupt binding. Consistent with this idea, introduction of R244E in silico significantly reduced the binding likelihood of TolB to CL-containing membranes, to ~49% chance (Fig 3C). We further demonstrated that TolB^R244E^ no longer exhibits significant heat changes when titrated into liposomes derived from *E. coli* polar lipids (Fig 3D), indicating loss of its ability to bind CL. TolB^R244E^ also does not affect liposome sedimentation (Fig S5). Taken together, we conclude that CL binding via R244 is critical for TolB function as part of the Tol-Pal complex to maintain OM lipid homeostasis in wild-type cells.

### The requirement for TolB-CL binding for OM lipid homeostasis is partially bypassed in strains lacking CL

Since CL binding is important for TolB function, one might expect CL-deficient strains to exhibit phenotypes associated with an impaired Tol-Pal complex. However, we found that Δ*clsABC* mutations (Δ*cls*) do not cause apparent OM permeability defects (Fig S6). These observations suggest that the presence of CL may not be strictly essential for Tol-Pal function in these cells. To further test this idea, we examined whether the CL binding ability of TolB is still required in the Δ*cls* mutant. TolB^R244E^ was produced at comparable levels to wild-type TolB in Δ*tolB* strains (Fig S7). To our surprise, TolB^R244E^ was able to partially rescue sensitivity of Δ*cls* Δ*tolB* strain to SDS-EDTA and vancomycin (Fig 4A). This mutation was even able to rescue the width defect in Δ*tolB* cells in the absence of CL (Fig 4B), overall demonstrating that TolB^R244E^ remains functional in OM lipid homeostasis in cells lacking CL. Collectively, our results suggest that although CL binding is important for TolB activity in wild-type cells, this requirement is partially bypassed when CL is absent, potentially due to compensatory changes in the membrane environment.

### TolB-CL binding is important for proper cell division

Despite having a primary function in maintaining OM lipid homeostasis, the Tol-Pal complex is recruited to mid-cell to facilitate OM invagination and/or septal cell wall separation. These roles during cell division are thought to involve Pal-mediated OM-cell wall tethering, modulation of cell wall enzymes, and/or possible OM lipid remodelling.^10, 18, 19^ We examined the impact of TolB^R244E^ on cell division phenotypes, and found that wild-type cells expressing this variant alone exhibited impaired growth and cell chaining under low osmolarity conditions (Fig S8). Interestingly, expressing TolB^R244E^ partially rescued growth defects, though not cell chaining in the Δ*cls* mutant. Using TolB-sfGFP constructs expressed at comparable levels (Fig S9), we showed that R244E did not affect enrichment of TolB to the mid cell in either wild-type or Δ*cls* strains, under normal osmolarity conditions (Fig 5A and B). Furthermore, Pal-mCherry still localized to the cell septum in the presence of TolB^R244E^ regardless of whether CL was available, albeit less strongly than in strains expressing wild-type TolB (Fig 5C and D). Therefore, Pal-mediated OM-cell wall tethering at the division site is still significant when TolB^R244E^ is present, as opposed to no tethering in the absence of TolB. We conclude that loss of TolB-CL binding may impact cell division instead via its effect on OM lipid homeostasis.

**Fig 5:**
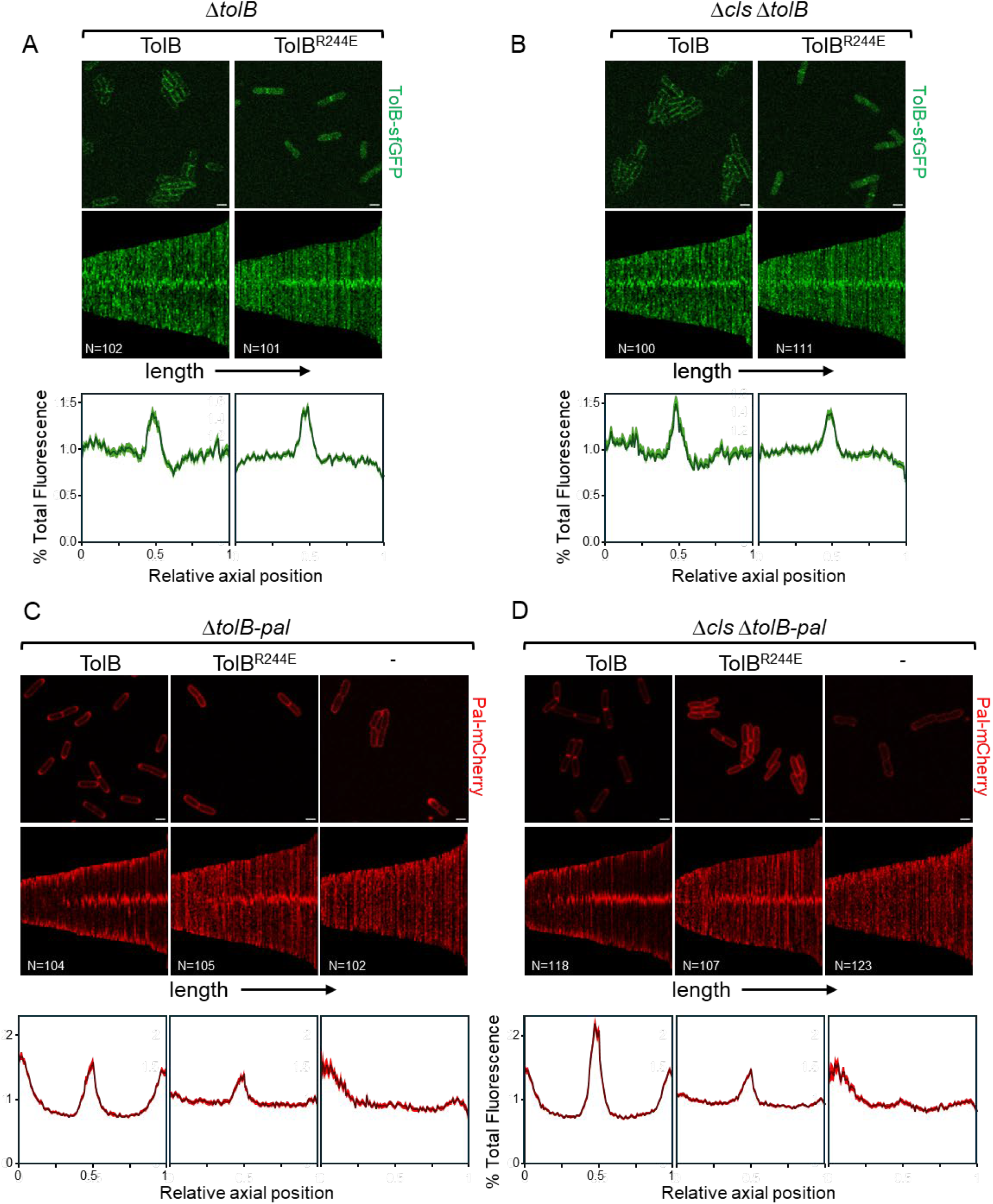
TolB and Pal still localize to cell septa in the presence of *tolB*^*R244E*^ mutation, and/or in the absence of CL. (*top*) Fluorescence microscopy images of (A) MG1655 Δ*tolB::kan* and (B). Δ*cls ΔtolB::kan* strains, expressing TolB-sfGFP or TolB^R244E^-sfGFP from pET22/42, and of (C). MG1655 Δ*tolB-pal::kan* and (D) Δ*cls ΔtolB-pal::kan* strains, expressing Pal-mCherry, TolB/Pal-mCherry or TolB^R244E^/Pal-mCherry from pBAD43. Scale bar represents 2 µm. (*middle*) Demographic representation of the fluorescence signals in the indicated strains measured along the long axis of individual cells, sorted according to cell length and aligned to mid-cell. (*bottom*) Average percentage distribution of the fluorescence signals along the long axis of all cells measured, where the relative axial position of 0.5 corresponds to mid-cell, and 0 to 1 correspond to the poles.

## Discussion

How the Tol-Pal complex maintains OM lipid homeostasis has remained elusive. While it has been implicated in retrograde transport of bulk PLs, there has been a lack of evidence for any protein-lipid interaction within the Tol-Pal complex.^10, 11^ In this study, we have uncovered the ability of the periplasmic protein, TolB, to bind lipid membranes in a CL-specific manner. We have shown that disruption of a putative CL-binding site on TolB abolished binding of CL-containing membranes, and impaired overall cellular function in OM lipid homeostasis. Interestingly, CL-binding became somewhat dispensable for Tol-Pal function in cells lacking CL, suggesting possible compensatory effects of other membrane lipids only when CL is absent. Overall, our work provides critical insights into the direct involvement of the Tol-Pal complex in lipid transport.

The discovery of TolB-CL interaction facilitates new ideas on how the Tol-Pal complex may maintain OM lipid homeostasis and transport. Due to its cone-shaped structure, CL is generally believed to be a non-bilayer lipid, which may preferentially localize to membrane regions of high (negative) curvature.^30^ This property is often attributed for why CL is enriched at the cell poles,^31^ albeit along with PG, a cylindrical bilayer lipid. In this vein, it is possible that CL would be found in other highly curved or strained membrane regions around the cell periphery. In cells lacking the Tol-Pal complex, excess PLs accumulate in the OM,^11^ which may induce local “ruffling” of the membrane, particularly in the inner leaflet. We speculate that CL may localize to such bulges in the OM, presumably of high curvature, therefore marking sites of excess lipids for TolB recognition. This process could be aided by Pal binding to TolB, positioning TolB closer to the inner leaflet of the OM. Binding of TolB to CL at these sites may facilitate subsequent interactions with TolA, which exerts a pulling force via the TolQR motor-stator complex at the IM. Therefore, CL may serve as a spatial cue for TolB recruitment, enabling the Tol-Pal complex to possibly mediate lipid transport, and maintain OM homeostasis. It is likely that CL is the dominant non-bilayer lipid that accumulates at sites of high curvature, explaining the requirement for TolB-CL binding in wild-type cells. In the absence of CL, and only so, we reason that other non-bilayer lipids, e.g. PA or PE, could then play compensatory role(s), rendering TolB-CL interactions non-essential. Further (biophysical) characterization of TolB-membrane binding will be critical. Given the essential role of TolB in OM homeostasis, its membrane-binding activity could be exploited as a potential drug target.

The primary function of the Tol-Pal complex is to maintain OM lipid homeostasis, possibly via retrograde PL transport.^10, 11^ In this context, given its ability to bind membranes via CL (head groups), TolB is quite likely the main effector. However, it is unlikely that TolB moves CL as a specific cargo, since cells lacking the Tol-Pal complex accumulate excess major PLs, not only CL, in the OM.^11^ Interestingly, the identified binding site to CL lies in the C-terminal β-propeller domain of TolB, lining the region(s) where TolB interacts with colicins or with Pal.^24, 32^ Instead, we propose that TolB binding to CL at membrane bulges may allow its N-terminal “TolA box” to become accessible (to TolA), similar to the effect of binding to colicins. The pulling force thus transduced by the TolQRA complex on TolB and its associated OM leaflet could bring this assembly closer to the IM, eventually leading to hemifusion, and possibly retrograde diffusion of PLs across a membrane bridge.^13^ Here, the role of Pal as a cell wall anchor may define permissive sites across the peptidoglycan mesh, and facilitate such trans-envelope membrane dynamics while preserving overall cell envelope integrity. This mechanistic working model, while speculative, provides a useful framework for our continued understanding of the Tol-Pal complex and its function in OM lipid homeostasis and transport in Gram-negative bacteria.

## Material and Methods

### Strains, plasmids, and growth conditions

The strains, plasmids and primers used were summarized in Table S1-4. All strains were grown in lysogeny broth (LB – 1% (w/v) tryptone and 0.5% (w/v) yeast extract with or without 1% (w/v) NaCl) liquid culture at 37 °C or on solid LB agar plates (1.5% (w/v) Bacto-Agar added to LB media) supplemented with ampicillin (100 µg/mL), spectinomycin (20 µg/mL), chloramphenicol (30 µg/mL), 1 mM Isopropyl β-D-1-thiogalactopyranoside (IPTG). The gene deletion alleles were constructed using λ Red recombination and transduced into strains using P1 transduction.^33^ Plasmids expressing TolB-sfGFP, Pal-mCherry, TolB-Pal-mCherry were constructed using Gibson assembly.^34^

### Isothermal Calorimetry (ITC)

TolB, sPal and Pal proteoliposomes in PBS were used for ITC. All proteins used in the experiment were in the same buffer (20 mM phosphate, 300 mM NaCl) to ensure no heat release from the mixture of different buffers. Prior to performing ITC, samples were centrifuged (10,000 × *g*, 4 °C, 5mins) to remove any precipitate. ITC was carried using NanoITC Microcalorimeter (TA Instruments). 50 μL of TolB at concentration of 140 μM (7 nmol) was injected into 20μM (166 μL, 3.32 nmol) of sPal or Pal proteoliposomes over 48 injections with an interval of 220 seconds. Experiments were carried out at 25 °C. 4.15 nmol of Pal proteins were used to generate Pal proteoliposomes, with ~80% surface-exposed (~3.32 nmol).

### Coarse-grained molecular dynamics simulations

Membrane binding of TolB was assayed using coarse-grained (CG) simulations. The monomeric TolB crystal structure was used as an input [Protein Data Bank ID code 2W8B; chain A.^24^ Protein atoms were converted to the CG Martini 3 force field^35^ using the martinize2 method.^36^ Additional bonds of 500 kJ mol^−1^ nm^−2^ were applied between all protein backbone beads within 1 nm. Proteins were oriented randomly using the Gromacs *editconf* tool, with a different starting orientation for each repeat. Proteins were placed 8 nm from a membrane (centre-of-mass distance), which were built using the insane protocol.^37^ Simulation boxes were 11 × 11 × 21 nm. All systems were solvated with Martini waters and Na^+^ and Cl^−^ ions to a neutral charge and 150 mM. Systems were minimized using the steepest descents method, followed by 1 ns equilibration with 5 fs time steps, then by 100 ns equilibration with 20 fs time steps, before 10 × 10 µs production simulations using 20 fs time steps, all in the NPT ensemble at 323 K with the V-rescale thermostat (τ_t_ = 1.0 ps) and semi-isotropic Parrinello-Rahman pressure coupling at 1 bar (τ_p_ = 12.0 ps). The Reaction-Field method was used to model long-range electrostatic interactions. Bond lengths were constrained to their equilibrium values using the LINCS algorithm. Simulations were run in Gromacs 2020.3.^38, 39^ Plots were made using Prism 10. MD snapshots were made using VMD.^40^ Data were analysed using Gromacs tools and the PyLipID package.^26^

### Efficiency of plating (EOP)

Strains were grown to mid-log (OD_600_ ~0.6) and normalized to an OD_600_ of 0.1. 10-fold dilution was performed to obtain a range of concentration of strains. 5µL of these diluted cultures were spotted on LB agar plates containing 0.5% sodium dodecyl sulfate (SDS)/ 0.5 mM Ethylenediaminetetraacetic acid (EDTA), 60 µg/mL vancomycin or without NaCl, unless otherwise stated. The LB agar plates were then incubated at 37 °C (for SDS EDTA and vancomycin plates) or at 42 °C (for plates without NaCl) for approximately 18hrs. All results shown are representative of at least three independent replicates.

### Microscopy

Overnight cultures of the strains were grown in LB and diluted (1:2000) in fresh media. The sub-cultures were grown to mid-log (~ 3 hrs) and concentrated 10-fold by centrifugation (3,000 × *g*, 5 mins). 10 µL of the sample was spotted on an agarose pad (1% agarose, in LB medium) and viewed under a confocal microscope (Olympus FV3000 Confocal Microscope). Images were analysed using ImageJ/FIJI (version 1.53c) and the MicrobeJ (version 5.13I) plugin.^41, 42^

## Acknowledgements

The authors would like to thank Jiang Hui Wu for her initial contributions in identifying residues for TolB-Pal crosslinking. We are grateful to Denis Duché (CNRS, Aix-Marseille Université) for providing the α-TolB and α-Pal antisera. We also thank Wee Boon Tan for useful and critical discussions. N.Z.-L.L. was supported by the National University of Singapore Graduate School of Integrative Sciences and Engineering Scholarship (ISEP). This work was supported by the Singapore Ministry of Health National Medical Research Council under its Open Fund Individual Research Grant (MOH-000145) and the Singapore Ministry of Education Academic Research Fund Tier 1 grant (National University of Singapore-Faculty of Science Preparatory Grant Scheme) (both to S.-S.C.). R.A.C. was funded by Wellcome (208361/Z/17/Z). P.J.S.’s lab was funded by Wellcome, MRC, BBSRC, EPSRC, NIH, JPIAMR and the Howard Dalton Centre. This project made use of time on ARCHER2 granted via the UK High-End Computing Consortium for Biomolecular Simulation, HECBioSim (http://www.hecbiosim.ac.uk), supported by EPSRC (grant no. EP/R029407/1). P.J.S. would like to thank the SCRTP at Warwick for use of the computing infrastructure. P.J.S. acknowledges Sulis at HPC Midlands+, which was funded by the EPSRC on grant EP/T022108/1.

The authors declare that they have no conflict of interest.

