## Supporting Information for "Lipid interactions are important for the Tol-Pal complex in maintaining outer membrane lipid homeostasis"

**Affiliations:**

This PDF file includes:

**Supplementary Material and Methods**

**Figures S1** to **S9**

**Tables S1** to **S4**

### Supplementary Material and Methods

#### Δ*clsABC* strain construction

*clsB* was first deleted from MG1655 strain via P1 transduction using BW25113 Δ*clsB*::*kan*. the resistance cassette marker was then removed using pCP20 plasmid and cured.[4] P1 transduction was used again to delete *clsC* gene from MG1655 Δ*clsB* strain using BW25113 Δ*clsC*::*cam*, and the resistance cassette marker removed using pCP20 and cured. Finally *clsA* was deleted from MG1655 Δ*clsBC* using BW25113 Δ*clsA*::*kan* and the resistance cassette marker was removed using pCP20 and cured.

#### Plasmid construction

pET22/42-*tolB-sfGFP* was cloned using Gibson assembly with the sfGFP gene sequence inserted into the pET22/42-*tolB* plasmid with the following junctions (different gene sequences are bolded):

| | | |
|---|---|---|
| pET22/42-*tolB*-*GGGGS* | 5'-TGGTCGCCGTATCTGGGTGGTGGTGGTTCA**TCTAAAGGTGAAGAA**-3' | ***sfGFP*** |
| *sfGFP* | 5'-GATGAGCTCTACAAACTCGAG**CACCACCAC**-3' | **pET22/42-*tolB*** |

pBAD43-*tolB-pal-mCherry* was cloned using Gibson assembly, the *tolB-pal-mCherry* operon from W3110 strain expressing chromosomal *pal-mCherry*, inclusive of 200 base pairs upstream of TolB was inserted into pBAD43 plasmid with the following junctions (different gene sequences are bolded):

| | | |
|---|---|---|
| pBAD43 | 5'-atcttctc**TTACTTGTACAGCTCGTCCATG**-3' | **Pal-mCherry** |
| Pal-mCherry | 5'-CAAGTAA**gagaagattttcagcctgatacagattaaatcag**-3' | **pBAD43** |

The *tolB* gene inclusive of the 200 base pairs upstream was then inserted into pBAD43-*pal-mCherry*

| | | |
|---|---|---|
| pBAD43 | 5'-ttaccaa**TCCCGAAACCACCAAGCC**-3' | **200bp upstream TolB** |
| 200bp upstream TolB | 5'-TTTCGGGA**ttggtaacgaatcagacaattgacg**-3' | **pBAD43** |

pBAD43-*pal-mCherry* was obtained by performing Gibson deletion to remove *tolB* from pBAD43-*tolB-pal-mCherry* (primer pair used are as listed below):

5'-CCAGATAAGGGAGATTAATAATTAATTGAATAGTAAAGGAATCATTGAAAT-3'

5'-ATCTCCCTTATCTGGACCGATGGCCCACGATAATTTA-3'

Plasmids pET22/42-*tolB-His*, pET22/42-*pal-His*, pET22/42-*spal-His* and pET22/42-*colA-His* were constructed using molecular cloning involving digestion and ligation. The target gene was amplified by polymerase chain reaction (PCR) using primers listed in **Table S3**.

Site-directed mutagenesis was performed using PCR with mutagenic primers listed in **Table S4**.

**Affinity purification**

BL21 (λDE3) cells were used to express proteins (Pal, soluble Pal (sPal), TolB, and Colicin A). Cells were grown in LB media and at mid-log phase ($OD_{600}$ ~0.8) 1 mM of IPTG was added to allow for protein expression for 2 hrs. Cells were harvested (4500 x *g*, 4 °C, 20 mins), and cell pellet was stored at -80 °C. Cell pellet was resuspended in 30 mL of phosphate-buffered saline (PBS), 300 mL NaCl buffer with 1 mM PMSF (calbiochem), 50 µg/mL DNase I (sigma), and 100 µg/mL lysozyme (calbiochem) added. French press (20,000 psi, French Press Glen Mills) was used to lyse the cells at high pressure. The lysed sample was centrifuged (4500 x *g*, 4 °C, 20 mins) to remove cell debris. Ultracentrifugation (90,000 x *g*, 4 °C, 1 hr) was used to separate the membrane and soluble fractions in the supernatant.

For soluble proteins (e.g., sPal, TolB, and Colicin A), the supernatant obtained after ultracentrifugation would be incubated with cobalt resins for 3 hrs to allow for isolation of protein through Co-TALON (Takara Bio) metal affinity purification. For membrane proteins (Pal), the pellet obtained after ultracentrifugation was resuspended in membrane solubilising buffer (PBS, 20 mM imidazole, 1% n-dodecyl-β-D-maltoside (DDM) (Avanti lipids)) and solution was incubated at 4 °C for 2 hrs. The mixture was then subjected to another round of ultracentrifugation (90,000 x *g*, 4 °C, 1hr) to separate the extracted membrane proteins from membrane debris. The supernatant obtained after ultracentrifugation would be incubated with cobalt resins for 3 hrs to allow for isolation of protein through Co-TALON metal affinity purification.

Following the incubation with cobalt resins, the solution was drained through gravity, and the filtrate was loaded back into the column and drained again. The column was washed 20x column volume of wash buffer (soluble proteins – PBS, 20 mM imidazole; membrane proteins – PBS, 20 mM imidazole and 0.05% DDM). After washing the cobalt resins, proteins were eluted with elution buffer (soluble proteins – PBS, 200 mM imidazole; Pal proteins – PBS, 200 mM imidazole, 0.05% DDM).

The eluted proteins were purified using size exclusion chromatography (SEC) (TolB – Superdex 200 increase 10/300 GL, Cytiva; Pal and sPal – Superdex 75 increase 10/300 GL, cytiva) was used with SEC running buffers (TolB and sPal – PBS, Pal – PBS, and 0.05% DDM). Purified proteins were flash frozen using dry ice and ethanol, then stored at -80 °C.

**Generating Pal proteoliposomes and assessing insertion orientation**

10 mg of *E. coli* polar lipids (Avanti lipids) were dissolved in 100 µL of chloroform and dried overnight in the fume hood. 1 mL of PBS was used to resuspend the lipids. The lipids were subjected to a series of freeze-thaw (11 times), to improve homogeneity of the lipids, followed by extrusion 21 times (Avanti lipids Mini Extruder) through a 200 nm membrane filter to obtain consistent size liposomes. 0.153% DDM ($R_{sat}$ concentration) was added to destabilize the lipid membrane of the liposomes to allow for insertion of proteins. Purified Pal (0.23 mg) were incubated with the liposomes at 4 °C for 2 hrs and the DDM was later removed by incubating twice with 100 mg methanol activated SM-2 Bio-beads.[1] These Pal proteoliposomes were directly used for ITC.

To determine the orientation of Pal in these liposomes. Proteoliposomes (0.2mg) were incubated with Proteinase K (100 µg/mL, at 25 °C) and at 10min timepoint aliquots of reaction mixture was quenched with 3 mM PMSF and 2x SDS reducing buffer. At 10min timepoint, $R_{sol}$ concentration of DDM (1% DDM) was added to the reaction to fully solubilize the liposome to allow exposure of internal Pal proteins to proteinase K. Reaction was quenched after 1min. Samples were heated for 10 mins at 100 °C and analysed using SDS-PAGE visualized with Coomassie blue staining and immunoblotting with antibodies.

**In vivo photoactivable crosslinking**

Site-directed mutagenesis was used to introduced amber stop codon (TAG) at position G396 in pET23/42-*tolB* plasmid. Overnight cultures of MG1655 Δ*tolB::kan* containing pSup-BpaRS-6TRN[2] and either pET23/42-$tolB^{G396x}$-*His* or pET23/42-$tolB^{G396x/R244E}$-*His* were diluted (1:100) in 1.5 L of fresh LB supplemented with 0.5mM *p*-benzoyl-L-phenylalanine (*p*Bpa) (GL shanghai) and grown to $OD_{600}$ ~ 1.0.[2, 3] Cells were harvested and affinity purification was performed. Samples were analysed using SDS-PAGE and immunoblotting with appropriate antibodies.

**Colicin spot assay**

Overnight cultures of the strains were grown in LB and diluted (1:100) in fresh media, supplemented with ampicillin and IPTG at 37 °C. The sub-cultures were grown to mid-log and normalized to $OD_{600}$ 0.005 by adding an appropriate volume of each culture to 10mL of molten soft LB agar (0.7% (w/w) Bacto-Agar in LB broth) containing ampicillin and IPTG. Each normalized soft agar culture was poured onto LB agar plate and allowed to cool and solidify. 10-fold dilution was performed to obtain a range of concentration of colicin A (100 mM, 10 mM, 1 mM, 100 nM, 10 nM, and 1 nM). 2 µL of each colicin A concentration was spotted onto the plates overlaid with normalized soft agar cultures. The plates were incubated at 37 °C overnight and inspected for inhibition zone on bacteria lawn. Images were captured by G:Box chemi-XT4 (Genesys version 1.3.4.0, Syngene).

**Sedimentation assay**

10 mg of lipids (*E. coli* polar lipids or synthetic lipids (3:1 POPE: POPG)) were dried overnight and hydrated with 1mL of PBS. The solution was subjected to a series of freeze-thaw (21 times) and extruded through a 200nm filter using the Avanti mini extruder (11 times). The liposomes were then diluted in 10 mL of PBS and ultracentrifuged (90,000 x *g*, 4 °C, 30 mins). The liposome pellets were resuspended to get a final concentration of 500nM. 50 µL of 0.05 mg of protein (LolB or TolB) was added to the 50 µL of liposomes. The solution was incubated with FM1-43 (1 µg) for 1 hr at room temperature and the fluorescence intensity of the mixture was measured (excitation 510 nm, emission 600 nm). The solution was then subjected to centrifugation (21,000 x *g*, 1 hr, 21 °C). The top 50 µL of the supernatant was harvested and had its fluorescence intensity measured.

Retention percentage = $\frac{Total\ FI - Supernatant\ Fi}{(Total\ FI + Supernatant\ FI) \div 2} \times 100$

**Western blot**

Samples were analysed with SDS-PAGE (polyacrylamide gel electrophoresis) using 12% polyacrylamide stacking gel. Gels were visualized through immunoblotting analysis by transferring gel onto polyvinylidene fluoride (PVDF) membranes (Immun-Blot 0.2 µm, Bio-rad) using the semi-dry electroblotting system (Trans-Blot Turbo Transfer System, Bio-rad). Membranes were blot using 1x casein blocking buffer (Sigma). Followed by either α-His (Qiagen) (1:5000) conjugated with horseradish peroxidase (HRP) or rabbit polyclonal primary α-TolA/α-TolB/α-Pal (1:1000) antisera and secondary α-rabbit antibodies (abcam). Blots were developed with Luminata Forte Western HRP substrate (Merck Milipore) and visualized via chemiluminescence using G:Box Chemi XT 4 (Genesys version 1.3.4.0, Syngene).

### Supplementary Figures

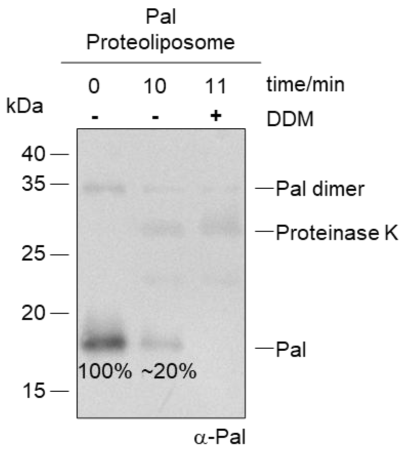


**Fig S1: Full length Pal is stably inserted into *E. coli* liposomes with a preferential outward-facing orientation.** SDS-PAGE/western blot analysis following limited proteinase K degradation of Pal reconstituted into liposomes comprising *E. coli* polar lipids. ~80% of Pal was digested after 10 min, indicating the majority of Pal was inserted on the exposed surface of the liposomes. The remaining ~20% of Pal facing the lumen became accessible to degradation upon the addition of the detergent DDM. Band intensities were measured using ImageJ, normalized to the 0-min timepoint.

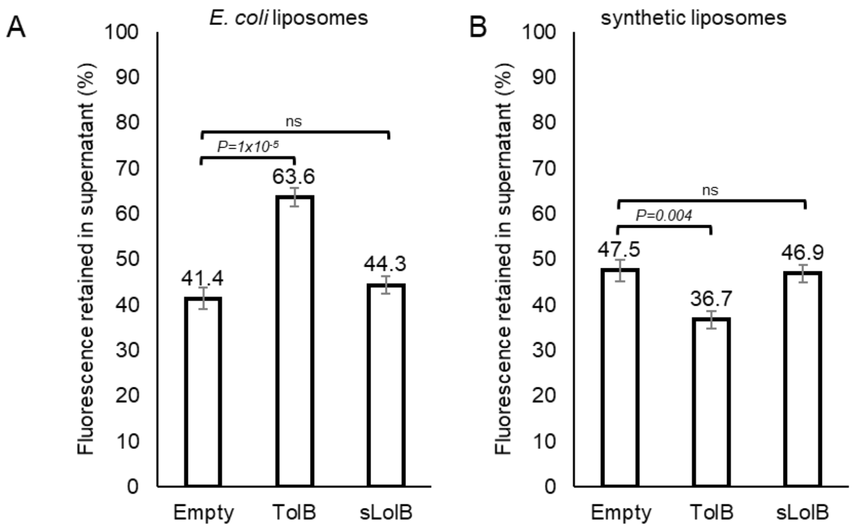


**Fig S2: TolB affects the sedimentation of liposomes comprising *E. coli* polar lipids.** (A) Representative sedimentation assays of (A) *E. coli* liposomes (PE:PG:CL 67:23:10), and (B) synthetic liposomes (PE:PG 3:1), in the presence of purified TolB or soluble LolB. FM1-43 dye was added to label the liposomes, and its fluorescence used as proxy for liposome concentration in the supernatant or pellet after 1-hr centrifugation. Technical triplicates were performed for each purified protein/ liposome preparation. Two-tailed Student's t-test was used to test for statistical significance.

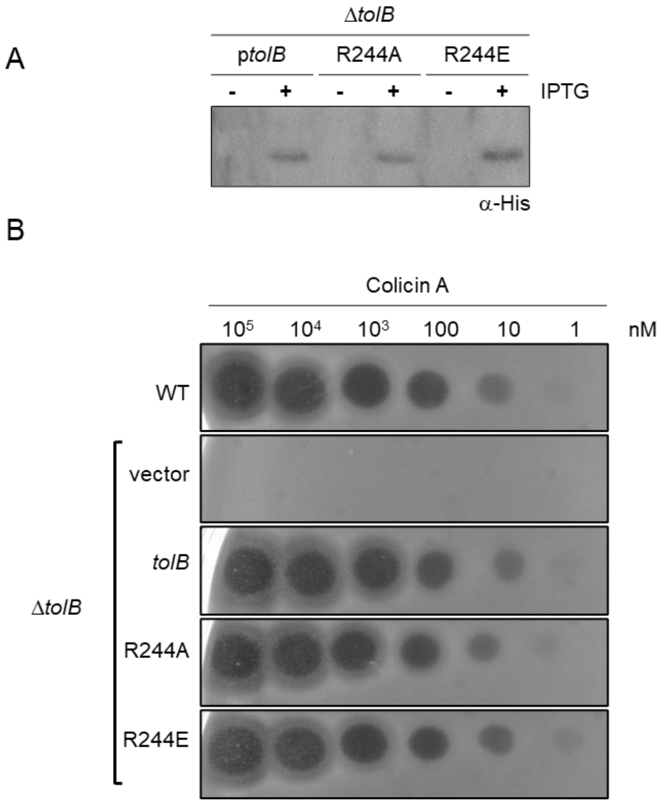


**Fig S3: The TolB$^{R244E}$ variant is stably expressed and fully functional in facilitating colicin entry.** (A) SDS-PAGE/western blot analysis of Δ*tolB* strain, containing pET22/42 expressing TolB-His and its R244 variants. 1 mM IPTG was added for de-repression of the *T7-lacO* promoter to facilitate leaky expression via endogenous RNA polymerases. (B) Spot test assays illustrating colicin A sensitivity of indicated strains on a lawn. Colicin A entry depends on TolB, and its interaction with the TolQRA complex.[5-7] TolB$^{R244E}$ retains its ability to interact with TolA and colicin A, implying structural stability.

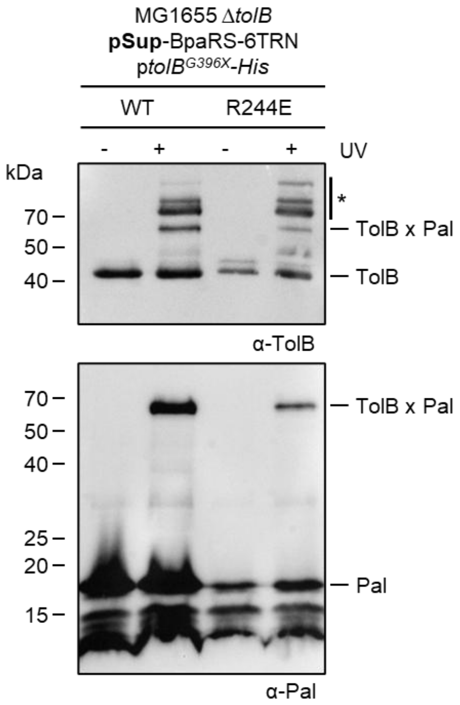


**Fig S4: TolB$^{R244E}$ interacts with Pal in cells at similar efficiency as TolB$^{WT}$.**

SDS-PAGE/Western blot analyses of affinity purification experiments using MG1655 Δ*tolB* strain containing either pET23/42-*tolB*$^{G396X}$-*His* or pET23/42-*tolB*$^{G396X/R244E}$-*His*, demonstrating non-covalent TolB-Pal interaction in the absence of UV irradiation and covalent TolBxPal crosslinks in the presence of UV irradiation. Introduction of unnatural amino acid *para*-benzoylphenylalanine (*p*Bpa) at a position 396 via amber codon suppression (G396X) enables UV-dependent crosslinking between TolB and Pal.[2] Based on band intensity quantification, efficiency of formation of crosslinks was comparable between TolB$^{WT}$ and TolB$^{R244E}$, where intensities of TolB$^{WT}$xPal and TolB$^{R244E}$xPal bands were ~40% and 33% of uncrosslinked TolB or Pal bands, respectively. Other yet-to-be-identified crosslinks to TolB are annotated with an asterisk (*).

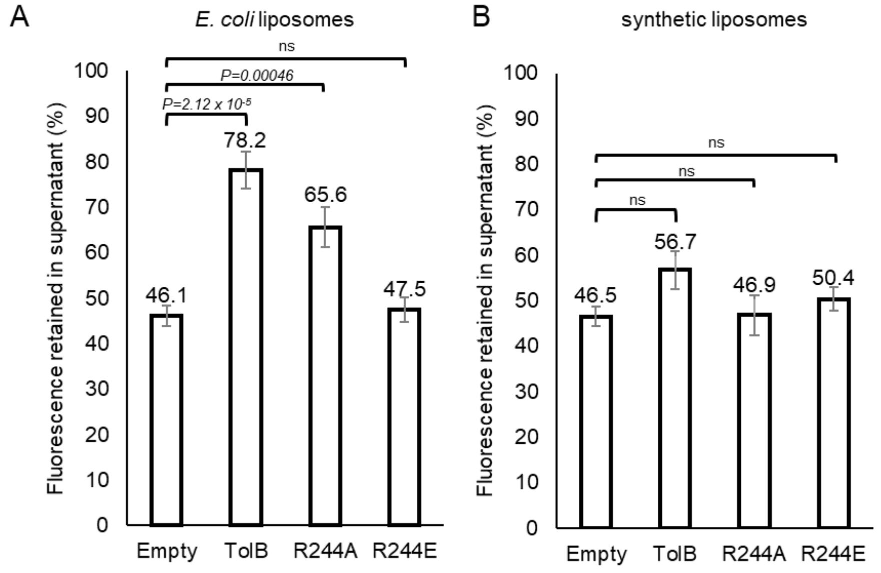


**Fig S5: TolB$^{R244E}$ no longer affects sedimentation of liposomes comprising *E. coli* polar lipids.** (A) Representative sedimentation assays of (A) *E. coli* liposomes (PE:PG:CL 67:23:10), and (B) synthetic liposomes (PE:PG 3:1), in the presence of purified TolB, functional TolB$^{R244A}$ or non-functional TolB$^{R244E}$ variants. FM1-43 dye was added to label the liposomes, and its fluorescence used as proxy for liposome concentration in the supernatant or pellet after 1-hr centrifugation. Technical triplicates were performed for each purified protein/ liposome preparation. Two-tailed Student's t-test was used to test for statistical significance.

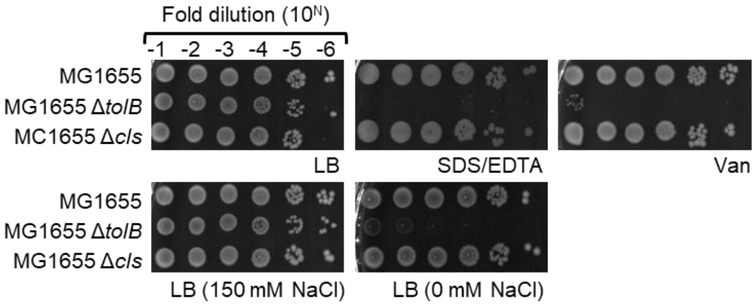


**Fig S6: Strains lacking CL do not exhibit *tol-pal* null phenotypes.** (A) EOP of MG1655, Δ*tolB*, and Δ*cls* strains, on LB agar plates supplemented with SDS/EDTA (0.5% SDS/0.5 mM EDTA), vancomycin (60 µg/mL) at 37 °C, or without NaCl at 42 °C.

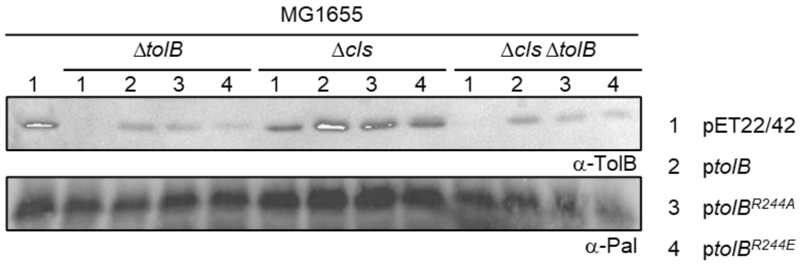


**Fig S7: TolB and its R244 variants are expressed at comparable levels within each background strain.** SDS-PAGE/western blot analyses of whole-cell lysates from MG1655 Δ*tolB*, Δ*cls*, and Δ*cls* Δ*tolB* strains, containing either pET22/42, pET22/42-*tolB*, pET22/42-*tolB*$^{R244A}$*,* or pET22/42-*tolB*$^{R244E}$. 1 mM IPTG was added for de-repression of the *T7-lacO* promoter to facilitate leaky expression via endogenous RNA polymerases.

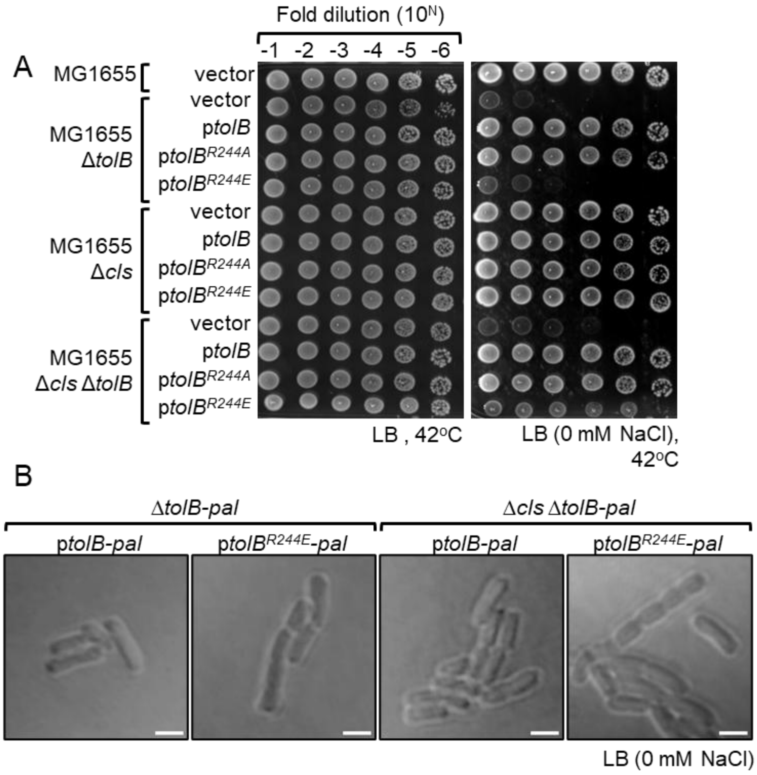


**Fig S8: TolB$^{R244E}$ partially rescues growth defects, but not cell chaining, in strains without CL.** (A) EOP of EOP of MG1655 wild type (WT), Δ*tolB*, Δ*cls*, and Δ*cls* Δ*tolB* strains, containing either pET22/42 empty vector or the plasmid expressing TolB and its mutants, on LB agar plates with and without NaCl at 42 °C. (B) DIC microscope images of MG1655 Δ*tolB-pal*, Δ*cls* Δ*tolB-pal* and Δ*tol-pal* strains, containing pBAD43-*tolB-pal-mCherry*, or pBAD43-*tolB$^{R244E}$-pal-mCherry*. Scale bar represent 2 μm. Cropped images are representative of at least three different uncropped field-of-view.

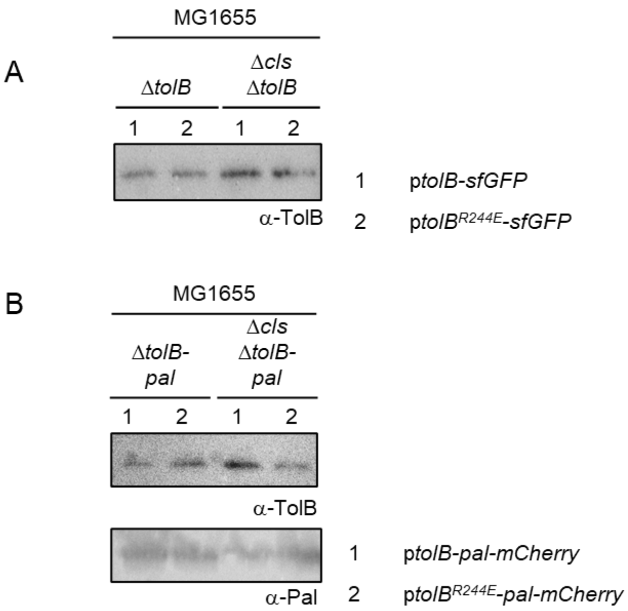


**Fig 9: Fluorescently-tagged TolB, TolB$^{R244E}$, and Pal can be stably expressed in different strains**. (A) SDS-PAGE/western blot analysis of whole-cell lysates from MG1655 Δ*tolB* and Δ*cls* Δ*tolB* strains, containing pET22/42-*tolB-sfGFP* or pET22/42-*tolB*$^{R244E}$-*sfGFP*. 1 mM IPTG was added for de-repression of the *T7-lacO* promoter to facilitate leaky expression via endogenous RNA polymerases. (B) SDS-PAGE/western blot analyses of whole-cell lysates from MG1655 Δ*tolB-pal* and Δ*cls* Δ*tolB-pal* strains, containing pBAD43-*tolB-pal-mCherry* or pBAD43-*tolB*$^{R244E}$-*pal-mCherry*.

**Table S1**

| Strain | Reference |
| --- | --- |
| MG1655 | Lab collection |
| MG1655 Δ*tolB*::*kan* | Lab collection |
| MG1655 Δ*clsABC* (labelled as Δ*cls*) | This study |
| MG1655 Δ*clsABC* Δ*tolB*::*kan* | This study |
| MG1655 Δ*tolB-pal*::*kan* | This study |
| MG1655 Δ*clsABC* Δ*tolB-pal*::*kan* | This study |
| MG1655 Δ*tol-pal*::*kan* | 8 |

Table S2

| Plasmids | Relevant genotypes and characteristics | Reference |
|---|---|---|
| pBAD43 | pBAD promoter, tight, arabinose-inducible expression, pSC101 ori, low-copy number plasmid | 9 |
| pET22/42 | pT7(lacO) inducible expression vector (enables leaky expression via endogenous RNA polymerases, when IPTG is added to de-repress lacO), contains multiple cloning site of pET42a(+) in pET22b(+) backbone; $Amp^R$ | 10 |
| pET23/42 | pT7 inducible expression vector lenables leaky transcription of T7 promoter by endogenous RNA), contains multiple cloning site of pET42a(+) in pET23a(+) backbone; $Amp^R$ | 11 |
| pET23/42-*tonB*$^{TM}$-*tolA* | Encodes the TonB$^{TM}$-TolA construct contains the first 34 amino acids from the N-terminus of TonB, including its transmembrane helix, fused directly to the periplasmic domains of TolA (amino acid 35-421) | 8 |
| pBAD33-*exbBD* | Encodes full length ExbBD | 8 |
| pBAD43-*tolB-pal* | Encodes full length TolB and Pal under control of *tolB* promoter | 8 |
| pSup-BpaRS-6TRN | Encodes an orthogonal tRNA and aminoacyl-tRNA synthetase permitting ribosomal incorporation of *p*Bpa at TAG stop codons (pSup) | 10 |
| pET22/42-*tolB-His* | Encodes full length TolB with C-terminal His8 tag | This study |
| pET22/42-*pal-His* | Encodes full length Pal with C-terminal His8 tag | This study |
| pET22/42-s*pal-His* | Encodes truncated soluble Pal (a.a. 65-173) with C-terminal His8 tag | This study |
| pET22/42-s*lolB-His* | Encodes de-lipidated LolB (a.a. 23-207) with C-terminal His6 tag and *pelB* signal peptide | 12 |
| pET22/42-*colA-His* | Encodes full length Colicin A with C-terminal His8 tag | This study |
| pBAD43-*tolB-pal-mCherry* | Encodes full length TolB and Pal fused with mCherry (with a GGGGS linker), under control of *tolB* promoter | This study |
| pBAD43-*pal-mCherry* | Encodes full length Pal fused with mCherry (with a GGGGS linker), under control of *tolB* promoter | This study |
| pET22/42-*tolB-sfGFP* | Encodes full length TolB fused with sfGFP (with a GGGGS linker) | This study |

**Table S3** (Molecular Cloning)

| Primers | Sequence (5' to 3') |
|---|---|
| NdeItolB | ACTACATATGAAGCAGGCATTACGAGTAGCATTT |
| XhoItolB | ACCACTCGAGCAGATACGGCGACCAGGCA |
| NdeIpal | ACTACATATGCAACTGAACAAAGTGCTGAAAGGG |
| XhoIpal | ACCACTCGAGGTAAACCAGTACCGCACGAC |
| NdeIsolpal | ACTACATATGCTGCAGCAGAACAACATCGT |
| XhoIsolpal | ACCACTCGAGGTAAACCAGTACCGCACGAC |
| NdeIcolA | ACTACATATGCCTGGATTTAATTATGGTGG |
| XhoIcolA | ACCACTCGAGATGTGCAGGTCGGATTATTT |

**Table S4** (site directed mutagenesis)

| Primers | Sequence (5' to 3') |
|---|---|
| tolBR223Afwd | GACCTTCGAAAGCGGTGCATCCGCGCTGGTTATTC |
| tolBR223Arev | GAATAACCAGCGCGGATGCACCGCTTTCGAAGGTC |
| tolBR223Efwd | GACCTTCGAAAGCGGTGAATCCGCGCTGGTTATTC |
| tolBR223Erev | GAATAACCAGCGCGGATTCACCGCTTTCGAAGGTC |
| tolBR244Afwd | CTTCATTCCCGGCTCACAACGGTGC |
| tolBR244Arev | GCACCGTTGTGAGCCGGGAATGAAG |
| tolBR244Efwd | GTGGCTTCATTCCCGGAACACAACGGTGCACC |
| tolBR244Erev | GGTGCACCGTTGTGTTCCGGGAATGAAGCCAC |
| tolBK264Afwd | CATTCGCCTTGTCGGCAACCGGTAGTCTG |
| tolBK264Arev | CAGACTACCGGTTGCCGACAAGGCGAATG |
| tolBK264Efwd | CATTCGCCTTGTCGGAAACCGGTAGTCTG |
| tolBK264Erev | CAGACTACCGGTTTCCGACAAGGCGAATG |
| tolBR287Afwd | GGTGACTGATGGTGCCAGTAACAATACC |
| tolBR287Arev | GGTATTGTTACTGGCACCATCAGTCACC |
| tolBR287Efwd | CAGGTGACTGATGGTGAAAGTAACAATACCGAAC |
| tolBR287Erev | GTTCGGTATTGTTACTTTCACCATCAGTCACCTG |
